# Physical activity predicts population-level age-related differences in frontal white matter

**DOI:** 10.1101/311050

**Authors:** Juho M. Strömmer, Simon W. Davis, Richard N. Henson, Lorraine K. Tyler, Cam-CAN, Karen L. Campbell

## Abstract

Physical activity has positive effects on brain health and cognitive function throughout the lifespan. Thus far, few studies have examined the effects of physical activity on white matter (WM) microstructure and psychomotor speed within the same, population-based sample (critical if conclusions are to extend to the wider population). Here, using diffusion tensor imaging and a simple reaction time task within a relatively large population-derived sample (N = 399; 18–87 years) from the Cambridge Centre for Ageing and Neuroscience (Cam-CAN), we demonstrate that physical activity mediates the effect of age on white matter integrity, measured with fractional anisotropy. Higher self-reported daily physical activity was associated with greater preservation of WM in several frontal tracts, including the genu of corpus callosum, uncinate fasciculus, external capsule and anterior limb of the internal capsule. We also show that the age-related slowing is mediated by WM integrity in the genu. Our findings contribute to a growing body of work suggesting that a physically active lifestyle may protect against age-related structural disconnection and slowing.

## 1. Introduction

Ageing is associated with profound changes in brain structure, including grey matter atrophy and alterations in the integrity of white matter (WM). Microstructural changes in the intra-and extracellular components of WM occur throughout the ageing brain, but tend to be more pronounced in frontal associative tracts ^1,2^. These age-related changes are thought to be driven largely by changes in myelin, with axon fibres being relatively unaffected by age ^3^. Fractional anisotropy (FA), an index of microstructural white matter integrity that is sensitive to changes in cerebral myelin levels, as indexed by post mortem histology ^4^ declines progressively with age in healthy adults, especially in those white matter tracts that mature later in life, such as anterior parts of corpus callosum ^5^.

This loss of myelin integrity is considered one of the key mechanisms underlying normal age-related variability in cognitive performance ^1,6^, often surpassing grey matter volume estimates in the ability to account for age-related cognitive decline ^7^. Increased WM integrity is often positively correlated with better performance across a number of cognitive domains and, in some circumstances, WM integrity mediates age-related slowing of cognitive processing ^8–12^. In fact, a substantial portion of the age-related variance in cognitive processing speed has been shown to be attributable to decreases in frontal WM integrity ^3,13^. The role of WM structures like the genu of corpus callosum in mediating the effect of age on cognitive processing speed has now been replicated many times ^2,12^ and this relationship appears to be specific to processing speed and executive functioning, rather than other aspects of cognition (e.g. language, motor functioning) ^10^. Thus, maintenance of WM structural connectivity appears to be particularly critical for the prevention of general age-related slowing. However, despite the ubiquity and cognitive relevance of these patterns of change in cerebral white matter, the specific mediators explaining these effects—beyond chronological age itself—are unclear.

While several lifestyle factors likely contribute to the maintenance of WM integrity with age, one of the most robust predictors of WM health appears to be physical activity. High cardiorespiratory fitness and engagement in physical activity have been shown to have protective effects for WM integrity ^14–16^ and cognitive performance ^17–19^ in healthy older adults. Evidence from prospective studies also indicate that physical activity considerably reduces the risk of dementia and Alzheimer’s disease ^20^. Interestingly, Burzynska et al. ^23^ showed that not only engagement in physical activity, but also avoiding sedentary behaviour, is important for preserving WM microstructural integrity later in life, possibly via different pathways. Sedentary lifestyle is more likely to be associated with obesity and poor aerobic fitness, and is a leading cause of disease and disability ^22^, which in turn, are shown to be associated with lower WM integrity ^23^. Longitudinal data from aerobic exercise intervention programs in older adults show that the selective increases in fitness associated with aerobic exercise, but not low-intensity control interventions, predict increases in WM integrity in the prefrontal and temporal cerebrum ^24^ and increases WM volume in the anterior corpus callosum ^25^. As noted above, these brain regions are particularly vulnerable to the detrimental effects of age. Together, these studies emphasize the potential benefits of physical activity in preventing age-related white matter loss.

While several studies suggest a link between exercise and differences in WM integrity with age ^24,26^, it remains to be seen whether this relationship holds within a large, population-based lifespan sample. Population-based samples are critical if our conclusions are to extend beyond relatively select (and potentially biased) samples of research volunteers to the population in general. Moreover, few studies, if any, have examined the relationship between brain health and participants’ reports of everyday activities and routines (encompassing such activities as cleaning the house and mode of transportation/distance to work), which arguably offer a more ecologically valid counterpoint to standard intervention studies ^27^. In this study, we examined the relationship between age, self-reported physical activity, WM microstructure, and processing speed within a large, population-based sample from the Cambridge Centre for Ageing and Neuroscience (Cam-CAN) ^28^. Participants (N = 399) completed a physical activity questionnaire ^29^ and series of cognitive tests, including simple reaction time (RT) task in their homes, before undergoing a series of structural and functional MRI scans, which included diffusion tensor imaging, DTI ^28^. DTI was used to estimate fractional anisotropy (FA) within 21 major tracts from the John Hopkins University (JHU) White Matter Atlas and related to physical activity and processing speed separately in a series of mediation models.

Our first objective was to determine whether physical activity mediates age-related decline in WM within particular tracts, and whether these are the tracts that are most susceptible to age-related decline. To this end, separate mediation models were run for each tract, testing whether the relationship between age and FA was mediated by daily physical activity. Our second objective was to examine whether performance on the simple RT task is associated with WM integrity and whether the age-related decline in this measure is mediated by WM integrity. Only those tracts that showed a significant mediation effect of physical activity in the first model (corrected for multiple comparisons), were included into the second set of models testing the association between age, FA and RT. Thus, our planned analyses will help to elucidate a possible explanation for age-related declines in white matter health and provide evidence for the role of this measure in predicting declines in processing speed.

## 2. Methods

### 2.1 Subjects

A healthy, population-based sample of 708 participants (age range 18 – 88 years), was collected as part of the Cambridge Centre for Ageing and Neuroscience (Cam-CAN; for detailed description of the study, see Shafto et al., 2014). The ethical approval for the study was obtained from the Cambridgeshire 2 (now East of England - Cambridge Central) Research Ethics Committee. Participants gave written informed consent. Exclusion criteria included poor vision (below 20/50 on Snellen test ^32^), poor hearing (failing to hear 35dB at 1000Hz in either ear), low MMSE (24 or lower ^31^, self-reported substance abuse (assessed by the Drug Abuse Screening Test (DAST-20; Skinner, 1982), poor English knowledge (non-native or non-bilingual English speaker), current psychiatric disorder or neurological disease. Additionally, people with contraindications to MRI or MEG were excluded. Handedness was assessed using Edinburgh Handedness Inventory ^33^. Of the initial 708, 646 participants had valid T1, T2 and DTI/DKI data. We also excluded participants who did not complete the RT task (N = 75), and those with outlying FA values further than 3 times interquartile range above or below the age decile mean (N = 25; total remaining N = 399, 221 females, age range 18 to 87 years). The sample characteristics are described in Table 1.

**Table 1.**
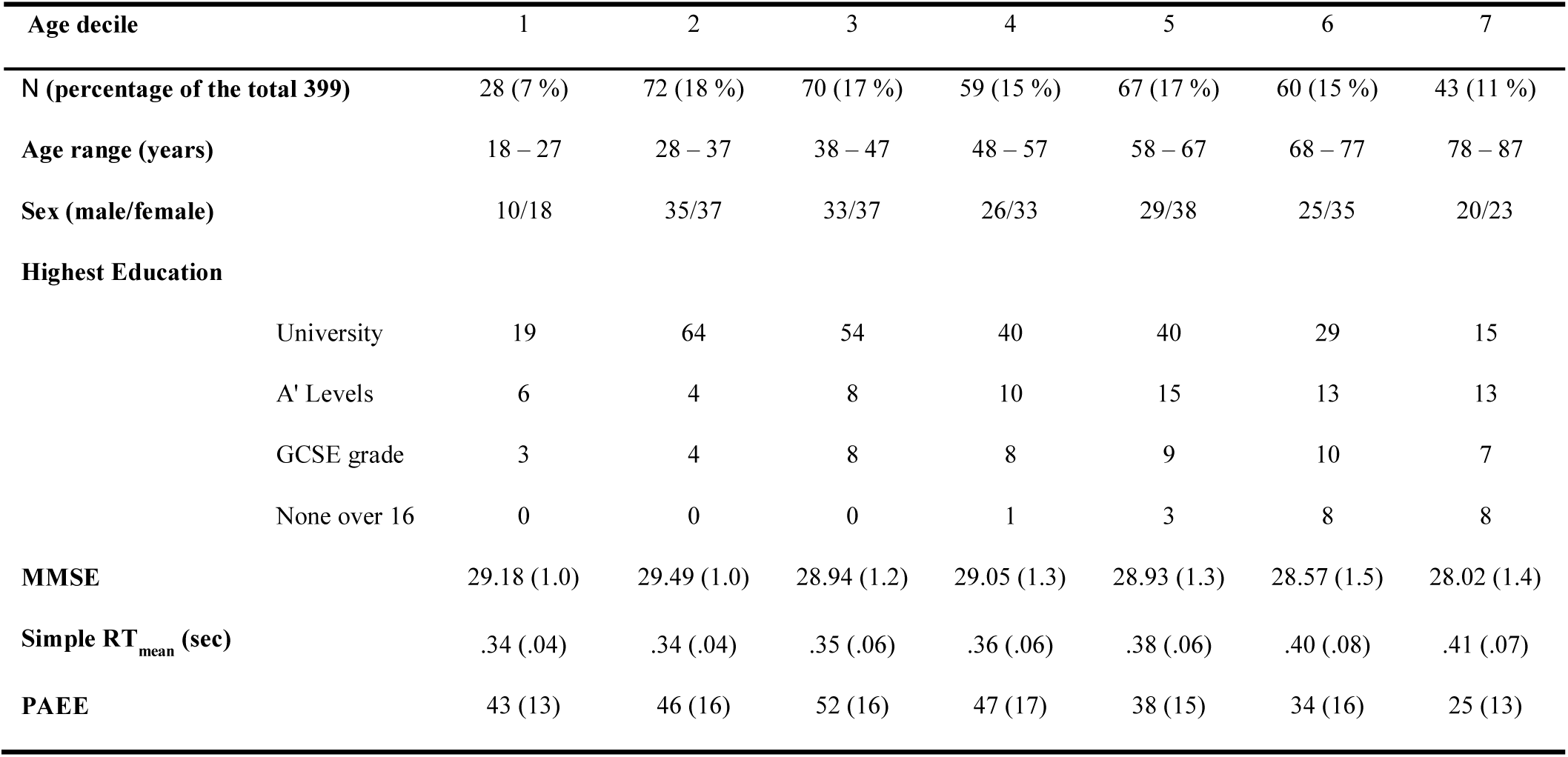
Participant demographic information. Values in parentheses are standard deviations. MMSE = mini mental status examination; Simple RT_mean_ = mean RT on the simple RT task. PAEE = physical activity energy expenditure (kJ/day/Kg)

### 2.2 Imaging pre-processing and region-wise analysis

The MRI data were collected from a Siemens 3T TIM TRIO (Siemens, Erlangen, Germany). To estimate white matter integrity (WMI), diffusion-weighted images were acquired with a twice-refocused-spin-echo sequence, with 30 diffusion gradient directions each for b-values 1,000 and 2,000 s mm ^−2^, and three images acquired using a b-value of 0 (TE = 104 ms, TR = 9.1 s, voxel size = 2 × 2 × 2 mm^3^, field of view (FOV) = 192 × 192 mm^2^, 66 axial slices, GRAPPA acceleration factor = 2).

All pre-processing was completed using a combination of functions from FSL version 4.1.8 (*bet, eddy, dtifit*, and *TBSS*) and custom MATLAB scripts. The diffusion data were pre-processed for eddy currents and subject motion using an affine registration model. After removal of non-brain tissue, a non-linear diffusion tensor model was fit to the DWI volumes. Non-linear fitting of the diffusion tensor provides a more accurate noise modelling than standard linear model fitting and enables various constraints on the diffusion tensor, such as positive definiteness. The tensor’s eigensystem was used to compute the fractional anisotropy (FA) at each voxel; FA maps were spatially normalized into a standard stereotactic space using tract-based spatial statistics ^34^. Images were then smoothed with a 6 mm full width at half maximum Gaussian kernel to address possible residual errors and inter-individual variability and to ensure the normality requirements of parametric statistics were met, and then masked with a binarised version of each participant’s FA map, such that voxels below an FA threshold of 0.35 were not considered for further analysis.

Next, the mean FA values over 21 bilaterally symmetrical tract ROIs from the JHU White Matter Atlas (http://cmrm.med.jhmi.edu/) were extracted for subsequent analysis: genu of corpus callosum, body of corpus callosum, splenium of corpus callosum, column and body of fornix, fornix (cres), cerebral peduncle, anterior limb of internal capsule, posterior limb of internal capsule, retrolenticular part of internal capsule, anterior corona radiata, superior corona radiate, posterior corona radiate, posterior thalamic radiation, sagittal stratum, external capsule, cingulate gyrus, hippocampus, superior longitudinal fasciculus, superior fronto-occipital fasciculus, uncinate fasciculus and tapetum.

### 2.3 Physical activity questionnaire

Information about physical activity energy expenditure (PAEE) was gathered as part of a larger self-completed questionnaire, which asked about education, training, travel, hobbies, and social activities. The questions about physical activity were based on items from the European Prospective Investigation into Cancer Study-Norfolk Physical Activity Questionnaire (EPIC-EPAQ2 ^29^. The full questionnaire is provided in Supplementary Information. Individual total PAEE per day (kJ/day/kg) was calculated from self-reported activities into metabolic equivalents (METs) ^35,36^, based on the standard definition of 1 MET as 3.5 ml O2 per min per kg (or 71 J/min/kg) based on the resting metabolic rate ^37^. In addition, PAEE was divided into subtypes in relation to the nature of the activity in order to investigate their contribution to total PAEE and age-related differences in it. Work PAEE includes all activities performed at work; Home PAEE includes home and housework-related activity; Leisure PAEE includes all voluntary leisure activity and exercise; and Commute PAEE includes commuting to work and other travel.

### 2.4 Response time task

In the simple RT task, participants were seated behind a computer screen and rested their right hand on a response box with four buttons (one for each finger). On the screen, they viewed an image of a hand with blank circles above each finger. Participants were instructed to press with the index finger as quickly as possible whenever the circle above the index finger in the image turned black. On pressing the button, or after maximum 3 seconds, the circle became blank again, and the variable inter-trial interval began. The inter-trial interval varied pseudo-randomly with positively skewed distribution, minimum 1.8 seconds, mean 3.7 seconds, median 3.9 seconds, and maximum 6.8 seconds. The task included 50 trials and mean RT was calculated for correct trials after applying a 3 SD trim to the data.

### 2.5 Statistical analysis

Pearson’s correlation coefficients (partialling out gender and education) were computed to examine the relationship between age and total PAEE. For the PAEE subtypes (which were skewed in their distributions), Spearman’s rank correlation coefficients were computed (partialling out gender and education) to examine age-related changes in the types of activity contributing to total PAEE.

In order to test whether physical activity helps to predict the effects of age-related WM decline, we ran a series of mediation analyses, in which a third mediator variable fully or partially accounts for the relationship between an independent predictor and dependent outcome variables ^38^. In each analysis, the independent factor was age, the dependent factor was one of the 21 white matter tracts (i.e., mean FA within a tract) and the mediator was the amount of physical activity (Figure 2D). For those tracks that showed a significant mediation effect, we went on to test the cognitive significance of that effect by examining the relationship between WM in those tracts and age-related slowing (Figure 2). To this end, we ran another set of mediation analyses using age as the independent factor, simple RT as the dependent factor and mean FA within each of the previously identified tracts as the mediator (Figure 2E). Direct effects of age on FA and RT were also included in these regressions. Statistical significance for mediation analyses is typically signified by a significant attenuation of the relationship (beta value) between predictor and outcome variables, denoted here by a 95% confidence interval (CI) for standardized regression coefficient that does not cross zero. All significance tests were two-tailed and False Discovery Rate (FDR) ^39^ at 0.05 was applied to protect against familywise Type I error.

**Figure 1.**
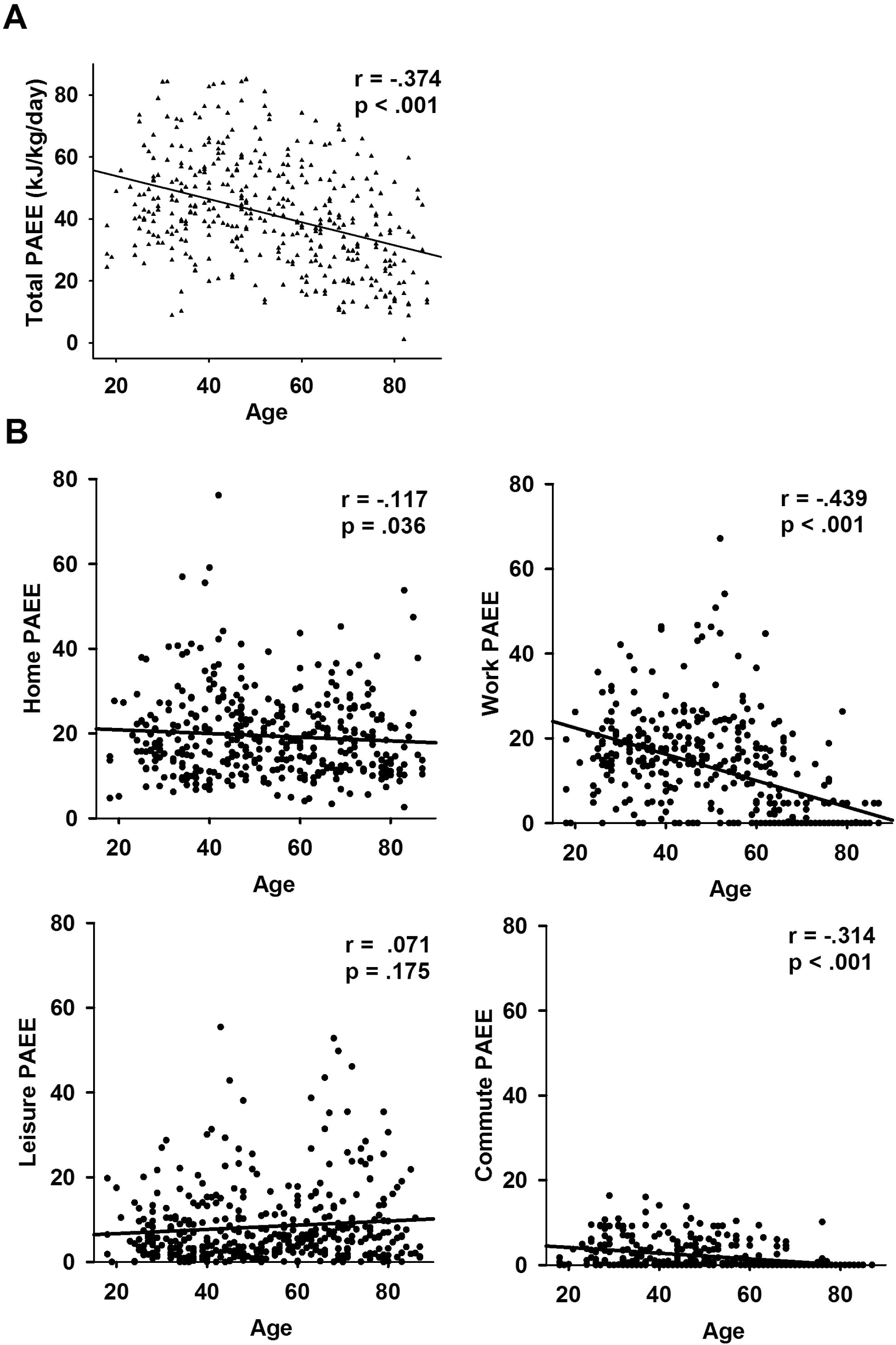
**A.** The effect of age on total physical activity energy expenditure (PAEE). **B.** The effect of age on PAEE subtypes of home-, work-, leisure-and commuting-related activities.

**Figure 2.**
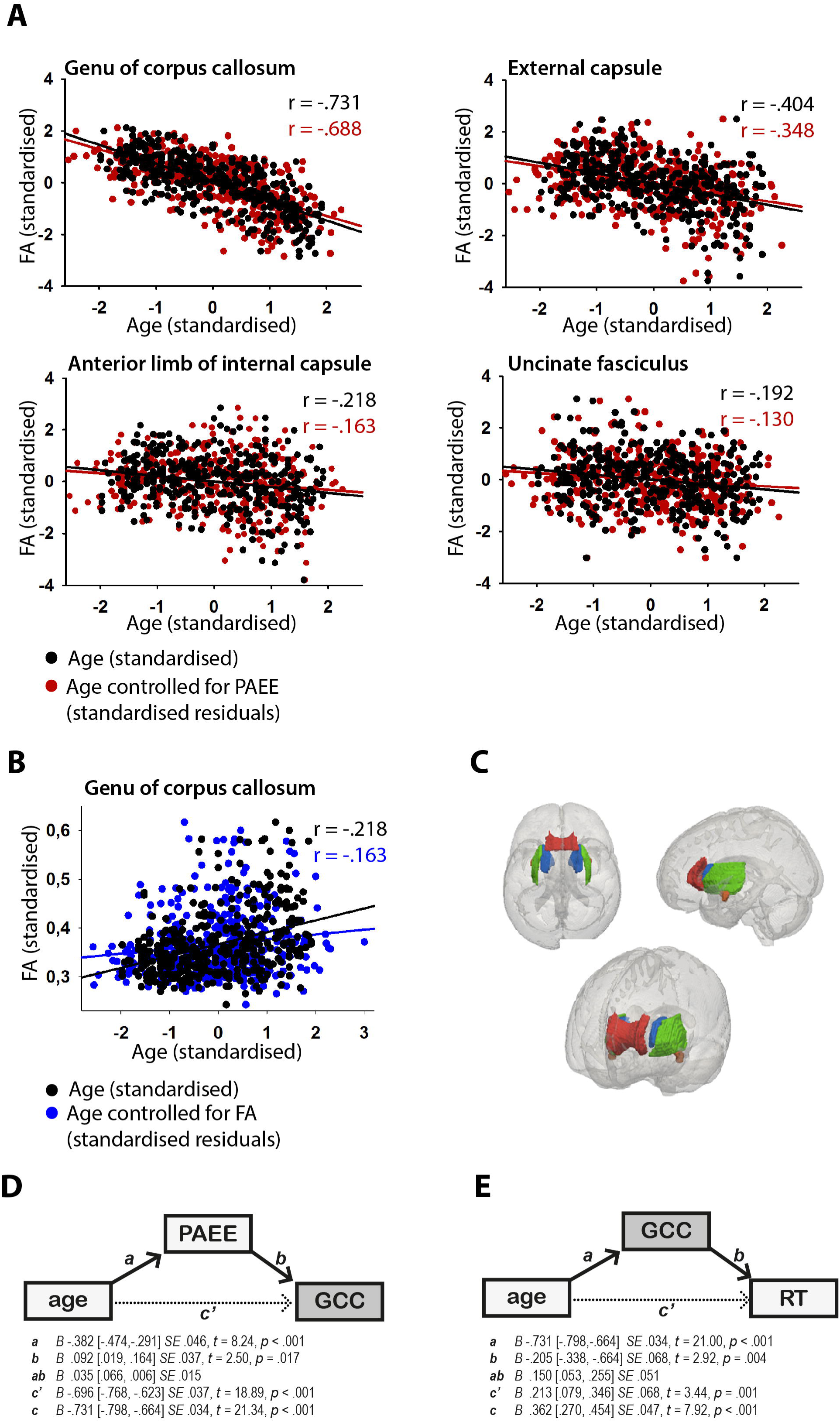
**A.** The relationship between white matter integrity (FA) and age (black dots) and age controlled for PAEE (red dots) in genu of corpus callosum, external capsule, anterior limb of internal capsule and uncinated fasciculus. FA decreases gradually with age within all of the analysed white matter tracts: GCC: r= -. 731, p < .001; EC: r= -. 404, p < .001; ALIC: r= -. 218, p < .001; UNC: r= -. 192, p < .001. The detrimental effect of age on FA is diminished in all of the analysed tracts when PAEE is partialled out from age: GCC: r= -. 688, p < .001; EC: r= -.348, p < .001; ALIC: r= -. 163, p = .001; UNC: r= -. 130, p = .009. The results indicate a positive relationship between higher physical activity and age-related differences in white matter microstructure. **B.** The relationship between reaction time and age (black dots) and age controlled for white matter integrity (FA) in genu of corpus callosum (red dots). Reaction times become gradually slower with age: r = .362, p < .001. The effect of age on reaction time is diminished when FA in genu of the corpus callosum is partialled out from age: r = .156, p = .002. The results indicate a positive relationship between white matter integrity in anterior corpus callosum and age-related differences in reaction time performance. **C.** White matter tract ROIs from JHU FA atlas. Tracts which survive the first stage of mediation analysis (genu, anterior limb of the internal capsule, and the external capsule) are rendered in (left to right) superior axial, sagittal, and oblique views. Genu – red; anterior limb of internal capsule – blue; external capsule – green; uncinate fasciculus – orange. **D.** Schematic representation of the mediation paths. PAEE mediates the effect of age on FA in genu of corpus callosum. **E.** FA in genu of corpus callosum mediates the effect of age on reaction time.

## 3. Results

### 3.1 Ageing and physical activity

Total PAEE, controlled for gender and education, showed a gradual decline with age: r = −0.37, p _fdr_ < .001 (Figure 1A). This is also shown in the results of the mediation models as path *a*, .i.e., the direct negative effect of age on total PAEE (Table 2). Work-related activity (rho = −0.52, p_fdr_< .001) and commuting-related activity (rho = −0.46, p_fdr_ < .001) showed moderate negative correlations with increasing age, but home-related activity showed a very weak correlation (rho = -.099, p _fdr_ = .06) and leisure time activity no correlation (rho = −0.09, p _fdr_ = .09) with age (Figure 1B). To conclude, leisure and home related activity seem to remain stable across the lifespan, while work and commuting-related activity decline and likely contribute to the decline in total PAEE.

**Table 2.** Mediation models testing the mediation of the relationship between age and white matter integrity (FA) in genu of corpus callosum, anterior limp of internal capsule, external capsule and uncinated fasciculus by physical activity energy expenditure (PAEE). *B* = standardized regression coefficient; SE = standard error. Asterisks denote significant mediation effects (for all effects, significance is denoted by a 95% confidence interval [CI] that does not cross zero; False Discovery Rate corrected *p* values < 0.05.).

| White matter tract | Path *a*<br>(age → PAEE) |  |  |  | Path *b*<br>(PAEE → FA) |  |  |  | Path *ab*<br>(mediation effect) |  | Path *c'*<br>(residual age → FA) |  |  |  | Path *c*<br>(age → FA) |  |  |  |
| --- | --- | --- | --- | --- | --- | --- | --- | --- | --- | --- | --- | --- | --- | --- | --- | --- | --- | --- |
|  | *B* [95% CI] | *B* SE | *t* | *p* | *B* [95% CI] | *B* SE | *t* | *p* | *B* [95% CI] | *B* SE | *B* [95% CI] | *B* SE | *t* | *p* | *B* [95% CI] | *B* SE | *t* | *p* |
| Genu of corpus callosum | -.382 [-.474, -.291] | .046 | 8.24 | <.001 | .092 [.019, .164] | .037 | 2.50 | .017 | -.035 [-.066, -.006]* | .015 | -.696 [-.768, -.623] | .037 | 18.89 | <.001 | -.731 [-.798, -.664] | .034 | 21.34 | <.001 |
| Anterior limb of internal capsule | -.382 [-.474, -.291] | .046 | 8.24 | <.001 | .118 [.014, .221] | .053 | 2.23 | .033 | -.045 [-.093, -.004]* | .023 | -.173 [-.277, -.070] | .053 | 3.29 | .001 | -.218 [-.315, -.122] | .049 | 4.46 | <.001 |
| External capsule | -.382 [-.474, -.291] | .046 | 8.24 | <.001 | .102 [.005, .200] | .050 | 2.07 | .049 | -.040 [-.083, -.003]* | .020 | -.365 [-.463, -.268] | .050 | 7.82 | <.001 | -.404 [-.495, -.314] | .046 | 8.81 | <.001 |
| Uncinate fasciculus | -.382 [-.474, -.291] | .046 | 8.24 | <.001 | .141 [.037, .245] | .053 | 2.66 | .011 | -.054 [-.100, -.013]* | .022 | -.138 [-.242, -.034] | .053 | 2.61 | .012 | -.192 [-.289, -.096] | .049 | 3.91 | <.001 |

### 3.2 Ageing and white matter integrity

The direct effect of age on FA was negative in all of the analysed tracts, except the posterior limb of internal capsule, which showed a small age-related increase in FA (Table 2, path *c*). The effect of age on FA was relatively large (standardized betas [*β’s*] < -.5) in the genu and body of the corpus callosum, fornix, anterior corona radiata, posterior thalamic radiation, sagittal stratum and tapetum (Figure 2A and 2C).

### 3.3 Physical activity and white matter integrity

The first mediation analyses tested whether total PAEE mediated the age-FA relationships. Four tracts showed a mediation effect that survived FDR correction: genu of corpus callosum, anterior limb of internal capsule, external capsule and uncinate fasciculus (Table 2, path *ab*, Figure 2A and 2C). The mediation effects of PAEE on these WM tracts are positive (Table 2, path *ab*), suggesting that higher physical activity is associated with less age-related white matter degeneration (see Figure 2A).^I^ No mediation effects were found when different PAEE types were used as mediator instead of total PAEE.

### 3.4 White matter integrity and speed of processing

The second mediation analyses tested whether FA (in the tracts related to exercise) mediated the relationship between age and processing speed. As expected, age was associated with slower responding on the simple RT task: *B =* .362, CI = .273, .452 SE = .047 (Table 3, path *c*). Critically, mean FA in the genu of corpus callosum (GCC) significantly mediated the effect of age on RT (ab = .150, CI = .045, .251, SE = .050; Table 3, path *ab*, Figure 2B), suggesting that preservation of white matter in the GCC is associated with less age-related slowing (Figure 2B). None of the other tracts showed significant mediation or main effects (Table 3, path *ab* and *c*).^II^

**Table 3.**
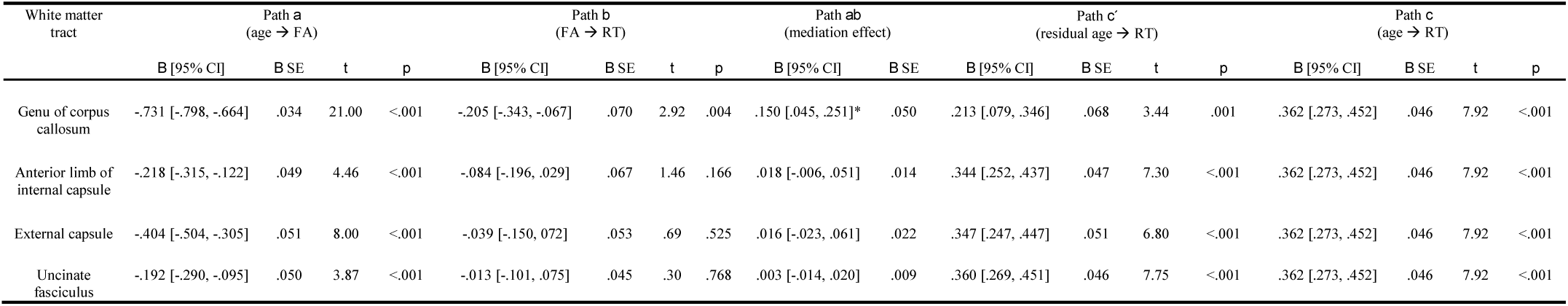
Mediation models testing the mediation between age and simple reaction time (RT), by correlation to the white matter integrity (FA) in genu of corpus callosum, anterior limp of internal capsule, external capsule and uncinated fasciculus. *B* = standardized regression coefficient; SE = standard error. Asterisks denote significant mediation effects (for all effects, significance is denoted by a 95% confidence interval [CI] that does not cross zero; False Discovery Rate corrected *p* values < 0.05.

## 4. Discussion

This study had two major aims. First, we examined whether physical activity mediates the effects of age on WM integrity. In line with previous work, we found higher physical activity to have positive effects that may protect against the damaging effects of age on FA in anterior WM tracts, namely the genu of corpus callosum, uncinate fasciculus, anterior limb of internal capsule and external capsule. The second aim of this study was to examine whether WM integrity within the tracts that benefit from physical activity mediate age-related slowing of processing speed. Of the four tracts tested, only the genu of the corpus callosum mediated a significant portion of the variance between age and RT on a simple motor task.

This is the first study, to our knowledge, to show a relationship between self-reported everyday activities and FA in a population-derived sample. While our results rely on a cross-sectional sample, and thus cannot relate physical activity to rates of longitudinal change, these results suggest that those who are more physically active in their day-to-day lives also have more youth-like patterns of WM microstructure. This is consistent with previous studies focusing on healthy older individuals, which have linked higher self-reported physical activity to higher WM volume ^40^ and lesser WM atrophy ^15^. Objectively measured cardiorespiratory fitness has also been shown to be associated with FA in the cingulum ^23^ and large portion of the corpus callosum (Johnson et al., 2012) in older adults. A recent study with two large samples of older adults demonstrated that white matter tracts between prefrontal regions and medial temporal lobe are particularly associated with cardiorespiratory fitness, and that these associations mediate spatial working memory performance ^41^. In our sample, which covers the whole adult age-range from 18 to 87 years, higher everyday physical activity was associated with less age-related loss of WM in several adjacent anterior tracts. Similarly, a recent study showed that higher cardiorespiratory fitness, assessed with the maximum volume of oxygen uptake (peak VO_2_), is related to higher FA in several WM tracts in older adults ^42^. Their study found regional specificity in the sensitivity to cardiorespiratory fitness – including genu of corpus callosum as one of the responsive regions. As with the current results, they showed that not all WM tracts that decline with age are associated with cardiorespiratory fitness.

Overall, physical activity declined with increasing age. This appears to be due largely to a decrease in activity related to work and commuting, whereas home-and leisure-related activity remained relatively stable across the age span. These results are in line with a recent review concluding that in childhood, habituation to active lifestyle, like active travel or outdoor play, are important contributors to total daily physical activity, whereas in adulthood, life events have the greatest influence on physical activity behaviour ^43^. In the present data, a drop in work-related activity around 60 years of age coincides with the mean retirement age in our sample. Thus, it may be that people whose everyday activity is highly dependent on the activities associated with work show the greatest drop in the total activity compared to those with an active lifestyle outside of working life. Thus, it seems particularly important to promote physical leisure activities amongst retired older adults, possibly with the help of societal actions.

Age-related slowing of cognitive processing has been proposed to underlie age-related declines within various domains of cognition ^44^. In the current study, simple RT slowed gradually with increasing age, which is a common finding among various types of age-related effects on speed of processing ^45^. Age-related slowing in RT was mediated by FA in the genu of corpus callosum, but not in the other tracts that related to physical activity. These findings are in line with an earlier study suggesting that WM deterioration in the anterior part of the corpus callosum may contribute to general age-related slowing ^2^, though other studies have also related the splenium of corpus callosum and anterior limb of internal capsule ^46^ and more global white matter structure ^12,47^ to perceptual-motor speed. A recent study also showed that lower whole brain FA is linked to inefficient brain response to cognitive demands of locomotion ^48^.

We acknowledge that our results do not speak to causality, since mediation analyses based on cross-sectional data do not inevitably represent causal relationships between age, physical activity, WM integrity and RT. Nevertheless, we assume that age, an independent factor in both of the mediation models, cannot be changed by the influence of other factors, and further, that psychomotor speed (RT) is a result of nervous system functioning (WM integrity), rather than the other way round ^49^. However, the causal interaction between lifestyle factors (e.g., physical activity) and brain structure remains unclear: it is well known that environment and behaviour, including physical activity, can cause plastic changes in the brain, but at the same time, changes in brain structure and function are known to influence behaviour (i.e. willingness towards action demanding physical activity). Furthermore, the strength of such inferences, based on self-reported questionnaire data, are necessarily limited. While the reliability of such questionnaires is high ^50^, their absolute validity is moderate at best. Thus, observations in large samples such as ours must be validated with more time-intensive vascular measures, such as VO2 uptake and neuroimaging measures of cerebral perfusion. In addition, it is important to note that the Cam-CAM sample represents the population in the UK and thus, these results may not generalize to a non-Caucasian population.

To conclude, we found that self-reported levels of physical activity mediated age-related white matter loss in a number of anterior tracts. While bearing in mind the limitations of cross-sectional data and a mediation-based approach, our findings complement the evidence from previous work suggesting that a physically active lifestyle may have protective benefits against age-related structural disconnection and cognitive decline. The findings of this study further support public health recommendations about the benefits of leading a physically active lifestyle across the life span, including older adults.

## Author Contributions

The principal personnel, including Lorraine K. Tyler, of the Cam-CAN project designed the study. Simon W. Davis and Juho M. Strömmer analysed the data. The interpretation of data was done by Juho M. Strömmer, Karen Campbell and Simon W. Davis. The manuscript was prepared by Juho M. Strömmer and revised by all the co-authors of this study. All of the authors approve the final version of the manuscript to be published.

## Acknowledgements

We are grateful to the Cam-CAN respondents and their primary care teams in Cambridge for their participation in this study. We also thank colleagues at the MRC Cognition and Brain Sciences Unit MEG and MRI facilities for their assistance. The Cam-CAN corporate author consists of the project principal personnel: Lorraine K Tyler, Carol Brayne, Edward T Bullmore, Andrew C Calder, Rhodri Cusack, Tim Dalgleish, John Duncan, Richard N Henson, Fiona E Matthews, William D Marslen-Wilson, James B Rowe, Meredith A Shafto; Research Associates: Karen Campbell, Teresa Cheung, Simon Davis, Linda Geerligs, Rogier Kievit, Anna McCarrey, Abdur Mustafa, Darren Price, David Samu, Jason R Taylor, Matthias Treder, Kamen Tsvetanov, Janna van Belle, Nitin Williams; Research Assistants: Lauren Bates, Tina Emery, Sharon Erzinçlioglu, Andrew Gadie, Sofia Gerbase, Stanimira Georgieva, Claire Hanley, Beth Parkin, David Troy; Affiliated Personnel: Tibor Auer, Marta Correia, Lu Gao, Emma Green, Rafael Henriques; Research Interviewers: Jodie Allen, Gillian Amery, Liana Amunts, Anne Barcroft, Amanda Castle, Cheryl Dias, Jonathan Dowrick, Melissa Fair, Hayley Fisher, Anna Goulding, Adarsh Grewal, Geoff Hale, Andrew Hilton, Frances Johnson, Patricia Johnston, Thea Kavanagh-Williamson, Magdalena Kwasniewska, Alison McMinn, Kim Norman, Jessica Penrose, Fiona Roby, Diane Rowland, John Sargeant, Maggie Squire, Beth Stevens, Aldabra Stoddart, Cheryl Stone, Tracy Thompson, Ozlem Yazlik; and administrative staff: Dan Barnes, Marie Dixon, Jaya Hillman, Joanne Mitchell, Laura Villis.

## Footnotes

I These effects remain equal when controlling for gender and education, although gender has a direct effect on FA in anterior limb of internal capsule (*B* = -.426, SE = .096, 95% CI = -.616 – -.237). These analyses were also run with Body Mass Index (BMI) as a covariate in order to investigate the association between BMI and PAEE. There were no associations between BMI and PAEE, although BMI has a marginal direct effect on external capsule (*B* = -.100, SE = .051, 95% CI = -.200 – -.001).

II These effects remain equal when controlling for gender and education, although gender has a direct effect on FA in anterior limb of internal capsule (*B* = -.426, SE = .096, 95% CI = -.616 – -.237).

## References

1. Madden DJ, Bennett IJ, Burzynska A, Potter GG, Chen N, Song AW. Diffusion tensor imaging of cerebral white matter integrity in cognitive aging. Biochim Biophys Acta - Mol Basis Dis. 2012;1822(3):386–400. doi:10.1016/j.bbadis.2011.08.003.

2. Salami A, Eriksson J, Nilsson L-G, Nyberg L. Age-related white matter microstructural differences partly mediate age-related decline in processing speed but not cognition. Biochim Biophys Acta - Mol Basis Dis. 2012;1822(3):408–415. doi:10.1016/j.bbadis.2011.09.001.

3. Madden DJ, Spaniol J, Costello MC, et al. Cerebral white matter integrity mediates adult age differences in cognitive performance. J Cogn Neurosci. 2009;21:289–302.

4. Schmierer K, Wheeler-Kingshott CAM, Boulby PA, et al. Diffusion tensor imaging of post mortem multiple sclerosis brain. Neuroimage. 2007;35(2):467–477. doi:10.1016/j.neuroimage.2006.12.010.

5. Kochunov P, Williamson DE, Lancaster J, et al. Fractional anisotropy of water diffusion in cerebral white matter across the lifespan. Neurobiol Aging. 2012;33(1):9–20. doi:10.1016/j.neurobiolaging.2010.01.014.

6. Madden DJ, Bennett IJ, Song AW. Cerebral white matter integrity and cognitive aging: Contributions from diffusion tensor imaging. Neuropsychol Rev. 2009;19(4):415–435. doi:10.1007/s11065-009-9113-2.

7. Westlye LT, Walhovd KB, Dale AM, et al. Life-span changes of the human brain white matter: Diffusion tensor imaging (DTI) and volumetry. Cereb Cortex. 2010;20(9):2055–2068. doi:10.1093/cercor/bhp280.

8. Bennett IJ, Madden DJ. Disconnected aging: Cerebral white matter integrity and age-related differences in cognition. Neuroscience. 2014;276:187–205. doi:10.1016/j.neuroscience.2013.11.026.

9. Nilsson J, Thomas AJ, O’Brien JT, Gallagher P. White Matter and Cognitive Decline in Aging: A Focus on Processing Speed and Variability. J Int Neuropsychol Soc. 2014;20(03):262–267. doi:10.1017/S1355617713001458.

10. Gazes Y, Bowman FD, Razlighi QR, O’Shea D, Stern Y, Habeck C. White matter tract covariance patterns predict age-declining cognitive abilities. Neuroimage. 2015;125:53–60. doi:10.1016/j.neuroimage.2015.10.016.

11. Price D, Tyler LK, Neto Henriques R, et al. Age-related delay in visual and auditory evoked responses is mediated by white-and grey-matter differences. Nat Commun. 2017;8(May 2016):15671. doi:10.1038/ncomms15671.

12. Kievit RA, Davis SW, Griffiths JD, Correia MM, Henson RNA. A watershed model of individual differences in fluid intelligence. Neuropsychologia. 2016;91:188–198. doi:10.1101/041368.

13. Kochunov P, Coyle T, Lancaster J, et al. Processing speed is correlated with cerebral health markers in the frontal lobes as quantified by neuroimaging. Neuroimage. 2010;49(2):1190–1199. doi:10.1016/j.neuroimage.2009.09.052.

14. Johnson NF, Kim C, Clasey JL, Bailey A, Gold BT. Cardiorespiratory fitness is positively correlated with cerebral white matter integrity in healthy seniors. Neuroimage. 2012;59(2):1514–1523. doi:10.1016/j.neuroimage.2011.08.032.

15. Gow AJ, Bastin ME, Valde MC, et al. Neuroprotective lifestyles and the aging brain. Neurology. 2012.

16. Oberlin LE, Verstynen TD, Burzynska AZ, et al. White matter microstructure mediates the relationship between cardiorespiratory fitness and spatial working memory in older adults. Neuroimage. 2015;(131):91–101. doi:10.1016/j.neuroimage.2015.09.053.

17. Colcombe S, Kramer AF. Fitness effects on the cognitive function of older adults: A meta-analytic study. Psychol Sci a J Am Psychol Soc / APS. 2003;14(2):125–130.

18. Sofi F, Valecchi D, Bacci D, et al. Physical activity and risk of cognitive decline: A meta-analysis of prospective studies. J Intern Med. 2011;269(1):107–117. doi:10.1111/j.1365- 2796.2010.02281.x.

19. Sexton CE, Betts JF, Demnitz N, Dawes H, Ebmeier KP, Johansen-berg H. A systematic review of MRI studies examining the relationship between physical fitness and activity and the white matter of the ageing brain. Neuroimage. 2016;131:81–90. doi:10.1016/j.neuroimage.2015.09.071.

20. Hamer M, Chida Y. Physical activity and risk of neurodegenerative disease: a systematic review of prospective evidence. Psychol Med. 2009;39(1):3–11. doi:10.1017/S0033291708003681.

21. Burzynska AZ, Chaddock-Heyman L, Voss MW, et al. Physical Activity and Cardiorespiratory Fitness Are Beneficial for White Matter in Low-Fit Older Adults. PLoS One. 2014;9(9):e107413. doi:10.1371/journal.pone.0107413.

22. Bauer UE, Briss PA, Goodman RA, Bowman BA. Prevention of chronic disease in the 21st century: Elimination of the leading preventable causes of premature death and disability in the USA. Lancet. 2014;384(9937):45–52. doi:10.1016/S0140-6736(14)60648-6.

23. Marks B, Katz L, Styner M, Smith J. Aerobic fitness and obesity: relationship to cerebral white matter integrity in the brain of active and sedentary older adults. Br J Sports Med. 2011;45(15):1208–1215. doi:10.1136/bjsm.2009.068114.

24. Voss MW, Heo S, Prakash RS, et al. The influence of aerobic fitness on cerebral white matter integrity and cognitive function in older adults: Results of a one-year exercise intervention. Hum Brain Mapp. 2013;34(11):2972–2985. doi:10.1002/hbm.22119.

25. Colcombe SJ, Erickson KI, Scalf PE, et al. Aerobic exercise training increases brain volume in aging humans. Journals Gerontol Ser a-Biological Sci Med Sci. 2006;61(11):1166–1170. doi:61/11/1166.

26. Marks BL, Madden DJ, Bucur B, et al. Role of aerobic fitness and aging on cerebral white matter integrity. Ann N Y Acad Sci. 2007;1097:171–174. doi:10.1196/annals.1379.022.

27. Best JR, Rosano C, Aizenstein HJ, et al. Long-term changes in time spent walking and subsequent cognitive and structural brain changes in older adults. Neurobiol Aging. 2017;57:153–161. doi:10.1016/j.neurobiolaging.2017.05.023.

28. Shafto M a, Tyler LK, Dixon M, et al. The Cambridge Centre for Ageing and Neuroscience (Cam-CAN) study protocol: a cross-sectional, lifespan, multidisciplinary examination of healthy cognitive ageing. BMC Neurol. 2014;14(1):204. doi:10.1186/s12883-014-0204-1.

29. Wareham NJ, Jakes RW, Rennie KL, Mitchell J, Hennings S, Day NE. Validity and repeatability of the EPIC-Norfolk Physical Activity Questionnaire. Int J Epidemiol. 2002;31(1):168–174. doi:10.1093/ije/31.1.168.

30. Snellen H. Probebuchstaben Zur Bestimmung Der Sehscha?rfe. Utrecht, Van de Weijer; 1962.

31. Folstein MF, Folstein SE, McHugh PR. “Mini-mental state”. A practical method for grading the cognitive state of patients for the clinician. J Psychiatr Res. 1975;12(3):189–198. doi:0022-3956(75)90026-6 [pii].

32. Skinner HA. The drug abuse screening test. Addict Behav. 1982;7(4):363–371.

33. Oldfield RC. The assessment and analysis of handedness: the Edinburgh inventory. Neuropsychologia. 1971;9(1):97–113.

34. Peled S, Friman O, Jolesz F, Westin C-F. Geometrically constrained two-tensor model for crossing tracts in DWI. Magn Reson Imaging. 2006;24(9):1263–1270. doi:10.1016/j.mri.2006.07.009.

35. Ainsworth BE, Haskell WL, Herrmann SD, et al. 2011 Compendium of Physical Activities: a second update of codes and MET values. Med Sci Sports Exerc. 2011;43(8):1575–1581. doi:10.1249/MSS.0b013e31821ece12.

36. Ainsworth BE, Haskell WL, Leon AS, et al. Compendium of physical activities: classification of energy costs of human physical activities. Med Sci Sports Exerc. 1993;25(1):71–80. http://www.ncbi.nlm.nih.gov/pubmed/8292105. Accessed May 26, 2015.

37. Henry C. Basal metabolic rate studies in humans: measurement and development of new equations. Public Health Nutr. 2005;8(March):1133–1152. doi:10.1079/PHN2005801.

38. Baron RM, Kenny D a. The Moderator-Mediator Variable Distinction in Social The Moderator-Mediator Variable Distinction in Social Psychological Research: Conceptual, Strategic, and Statistical Considerations. J Pers Soc Psychol. 1986;51(6):1173–1182. doi:10.1037/0022-3514.51.6.1173.

39. Benjamini Y, Hochberg Y. Benjamini Y, Hochberg Y. Controlling the false discovery rate: a practical and powerful approach to multiple testing. J R Stat Soc B. 1995;57(1):289–300. doi:10.2307/2346101.

40. Benedict C, Brooks SJ, Kullberg J, et al. Association between physical activity and brain health in older adults. Neurobiol Aging. 2013;34(1):83–90. doi:10.1016/j.neurobiolaging.2012.04.013.

41. Oberlin LE, Verstynen TD, Burzynska AZ, et al. White matter microstructure mediates the relationship between cardiorespiratory fitness and spatial working memory in older adults. Neuroimage. 2015. doi:10.1016/j.neuroimage.2015.09.053.

42. Hayes SM, Salat DH, Forman DE, Sperling RA, Verfaellie M. Cardiorespiratory fitness is associated with white matter integrity in aging. Ann Clin Transl Neurol. 2015;2(6):688–698. doi:10.1002/acn3.204.

43. Condello G, Puggina A, Aleksovska K, et al. Behavioral determinants of physical activity across the life courselJ: a “ Determinants of Diet and Physical Activity “ (DEDIPAC) umbrella systematic literature review. Int J Behav Nutr Phys Act. 2017;14(58). doi:10.1186/s12966-017-0510-2.

44. Salthouse TA. The processing-speed theory of adult age differences in cognition. Psychol Rev. 1996;103(3):403–428. doi:10.1037/0033-295X.103.3.403.

45. Salthouse T a. Aging and measures of processing speed. Biol Psychol. 2000;54(1-3):35–54. doi:10.1016/S0301-0511(00)00052-1.

46. Madden DJ, Whiting WL, Huettel SA, White LE, Macfall JR, Provenzale JM. Diffusion tensor imaging of adult age differences in cerebral white matterlJ: relation to response time. 2004;21:1174–1181. doi:10.1016/j.neuroimage.2003.11.004.

47. Johnson M a, Diaz MT, Madden DJ. Global versus tract-specific components of cerebral white matter integrity: relation to adult age and perceptual-motor speed. Brain Struct Funct. 2014;(220):2705–2720. doi:10.1007/s00429-014-0822-9.

48. Lucas M, Wagshul ME, Izzetoglu M, Holtzer R. Moderating Effect of White Matter Integrity on Brain Activation During Dual-Task Walking in Older Adults. Journals Gerontol Ser A. 2018;XX(Xx):1–7. doi:10.1093/gerona/gly131.

49. Gow AJ, Corley J, Starr JM, Deary IJ. Reverse causation in activity-cognitive ability associations: The Lothian Birth Cohort 1936. Psychol Aging. 2012;27(1):250–255. doi:10.1037/a0024144.

50. Helmerhorst HJ, Brage S, Warren J, Besson H, Ekelund U. A systematic review of reliability and objective criterion-related validity of physical activity questionnaires. Int J Behav Nutr Phys Act. 2012;9(1):103. doi:10.1186/1479-5868-9-103.

